# Unravelling ant-symbiont network topology across Europe

**DOI:** 10.1101/859058

**Authors:** Thomas Parmentier, Frederik de Laender, Dries Bonte

## Abstract

Long-term associations between different species are key drivers in community composition in all ecosystems. Understanding the ecological and evolutionary drivers of these symbiotic associations is challenging because of the diversity of species and interaction types hosted in natural ecological networks. Here, we compiled the most complete database on natural ant-symbiont networks in Europe to identify the drivers of bipartite network topology. These ant-symbiont networks host an unrivalled diversity of symbiotic associations across the entire mutualism-antagonism continuum, of which the most diverse types of symbionts are (1) trophobionts: mutualistic aphids and scale insects (2) myrmecophiles: commensalistic and parasitic arthropods, and (3) social parasites: parasitic ant species. These diverse ant-symbiont networks provide a unique opportunity to tease apart ecological and evolutionary drivers. To do so, we dissected network topology and asked what determines host specificity and which host factors drive symbiont species richness and facilitate host switching for the different types of symbionts.

We found an unexpectedly high number of 701 obligate symbionts associated with European ants. Symbiont type explained host specificity and the average relatedness of the targeted host species. Social parasites were associated with few, but phylogenetically highly related hosts, whereas trophobionts and myrmecophiles interacted with a higher number of hosts across a wider taxonomic distribution. Colony size, host range and habitat type predicted total symbiont richness, where ants hosts with larger colony size or larger distribution range contained more symbiont species. However, we found that different sets of host factors affected diversity in the different types of symbionts. Ecological factors, such as colony size, host range and niche width predominantly drive myrmecophile species richness, whereas evolutionary factors, such as host phylogeny and biogeography, mainly determine richness of mutualistic trophobionts and social parasites. Lastly, we found that hosts with a common biogeographic history support a more similar community of symbionts. Phylogenetic related hosts also shared more trophobionts and social parasites, but not myrmecophiles. Taken together, these results suggest that ecological and evolutionary processes drive host specificity and symbiont richness in large-scale ant-symbiont networks, but these drivers may shift in importance depending on the type of symbiosis. Our findings highlight the potential of well-characterized bipartite networks composed of different types of symbioses to identify candidate processes driving community composition.

## I. INTRODUCTION

Long-term associations between different species, known as symbioses, are crucial components of community assembly in all ecosystems. These intricate associations display a high diversity, ranging from mutually beneficial partnerships to parasitic interactions, where one species exploits another (Paracer & Ahmadjian, 2000). Interactions among species are central to ecological and evolutionary dynamics in assemblages of species that belong to different guilds and trophic levels. They are as such essential elements of the entangled bank, Darwin’s metaphor for the complexity and connectedness of natural systems (Darwin, 1859), and give rise to important stabilising feedbacks that eventually maintain diversity and ecosystem functioning (Thrall *et al*., 2007; Bastolla *et al*., 2009). To date, these insights are merely derived from theory and empiricism focusing on either antagonistic competition and predator-prey interactions (e.g. Hairston, Smith, & Slobodkin, 1960; Pimm, 1979; Tilman, 1982) or more recently mutualistic interactions (Bascompte, Jordano, & Olesen, 2006; Bascompte & Jordano, 2007).

Communities contain a wide variety of interactions, rendering the ecological network very complex (Newman, 2003). One important feature of complex systems is the presence of emergent properties that result from interactions among the specific components of the system (Solé & Bascompte, 2006). Typically, the emergent properties result from the complex interaction between the different network components across time and space and are difficult to predict from the specific (isolated) pairwise interactions (Vázquez, Chacoff, & Cagnolo, 2009). Generally speaking, modular networks that are characterised by a high degree distribution and connectance tend to be more robust to species loss, and are less affected by disturbance (Solé & Bascompte, 2006; Olesen *et al*., 2007). While theoretical progress has been made (Solé & Montoya, 2001), the field is suffering from a lack of comprehensive and manageable empirical systems.

The representation of communities in networks constitutes a holistic approach, but the study and interpretation of the drivers and consequences of the topology will largely depend on the considered scale, both in terms of spatiotemporal and phylogenetic dimensions (Delmas *et al*., 2019). While network studies at local scales will be informative on the local community assembly processes, those reconstructed at regional or global scales will allow inference of macro-ecological and evolutionary processes by making abstraction of the formerly described local drivers (Trøjelsgaard & Olesen, 2013).

In contrast to trophic networks, bipartite host-symbiont networks contain different kinds of links, with interactions between hosts and their symbionts ranging from antagonistic to mutualistic (Ings *et al*., 2009). Examples of such networks are for instance plant-mycorrhiza and host-microbiome associations. The complexity of these networks is enormous, and their description is merely based on one interaction type, either antagonistic or mutualistic, although theory predicts that the diversity of interaction types may be essential for community stability (Fontaine *et al*., 2011;Mougi & Kondoh, 2012). The topology of bipartite host-symbiont networks can be dissected by adopting two different perspectives, i.e. that of each of the individual sets of species (hosts and symbionts). Asking what factors cause a given topology is equivalent to asking, for each of the sets, what explains the number of links per species and the specificity of these links, i.e. how links are distributed among species from the focal set. An example of this approach is found in studies on predator-prey networks, where average web vulnerability (i.e., the average number of predators per prey) and generality (i.e., the average number of prey eaten per predator) link the specificity of the two interacting species sets (Schoener, 1989).

A determining feature of the ecology of symbionts is host specificity which can be quantified in host-symbiont networks by the number of links departing from a symbiont node. Yet, a measure of host specificity should ideally consider the relatedness of the targeted host species as well (Poulin & Mouillot, 2003). Generalist symbionts target multiple, unrelated host species and are thought to gain relatively low benefits in any host. Specialist symbionts, in turn, engage with one or only a few related species, but achieve high benefits with their hosts relative to the generalists due to advanced morphological, physiological and behavioural adaptations (Bronstein, Alarcón, & Geber, 2006; Thrall *et al*., 2007). Their strong specialization, however, is offset by lower population densities and higher extinction risks due to the lower availability of hosts. Several studies in host-symbiont systems clearly found that host specificity is tightly linked with fundamental ecological processes and evolutionary history. Typically, host specificity is different among groups of the symbiont community as for example demonstrated in parasites of primates (Pedersen *et al*., 2005) and in parasitic mites on mussels (Edwards & Malcolm, 2016). A study on moths and plants indicated that host specificity can be dependent on the type of symbiotic interaction with pollinating moths being more specific than the parasitic leaf-feeding relatives (Kawakita *et al*., 2010).

From the perspective of the host, it is fundamental to understand the ecological, evolutionary and environmental drivers that promote the number of associated symbionts, i.e. the number of links departing from a host node to symbiont nodes. Studies on other host-symbiont systems reported multiple host variables which correlate with parasite species richness (PSR). Generally, the makeup of symbiont communities is orchestrated by both ecological and evolutionary host factors. The driving ecological factors are analogous to the ones that favour species richness on islands (cf. theory of island biogeography, MacArthur & Wilson, 1967) as host species can be regarded as biological islands (Kuris, Blaustein, & Alio, 1980). Symbiont richness is therefore expected to increase with the number and variety of microhabitats offered by the host, host longevity, host range, and the interaction probability with other hosts (Kamiya *et al*., 2014; Stephens *et al*., 2016). Evolutionary processes may affect symbiont species richness in different ways. Related hosts often show traits that are correlated throughout evolution (phylogenetic correlation) which lead to similar values in species richness. However, related host species may have diverged with time, whether or not in a common spatial evolutionary ancestry (biogeography), but may still attract a similar fauna of symbionts as unrelated host species with a similar ecology (Poulin, 1995).

Another pattern that emerges in host-symbiont networks is the sharing/transmission of symbiont species across host species. The degree of symbiont sharing is vital as symbiont transmissions can connect eco-evolutionary dynamics across hosts as a result from rapid symbiont spread in host populations (e.g. Jaenike *et al*., 2010; Himler *et al*., 2011 in endosymbionts). While little is known about the proximate mechanisms by which single symbionts switch between hosts, we can anticipate that host species with similar ecological niches and/or a shared evolutionary history tend to have similar symbiont communities. The pervasive effect of phylogenetic relatedness on symbiont sharing has for example been demonstrated in bat parasites (Luis *et al*., 2015) and in plant-mycorrhiza (Veresoglou & Rillig, 2014).

Ant-symbiont networks are ideally suited to study which factors drive bipartite network topology (Ivens *et al*., 2016). The diversity of symbiotic associations found in ants (Kistner, 1982; Hölldobler & Wilson, 1990; Rettenmeyer *et al*., 2010; Parmentier, in press) is thought to be promoted by their omnipresence in terrestrial ecosystems, their stable and climate-controlled nest fortresses and the high number of available resources in the nest (Kronauer & Pierce, 2011). Ants interact with different types of symbionts spanning the entire parasitism-mutualism gradient. They include parasitic ants, different groups of arthropods living in the nests, mutualistic aphids, nematodes, plants, bacteria and fungi. Therefore, they are ideal systems to study different interaction types in a single framework (Fontaine *et al*., 2011).

Ant-symbiont networks that have been recently studied, deal with local interaction networks and mostly focus on one kind of symbiotic interaction in isolation, such as mutualistic plant-ant networks (Guimaraes *et al*., 2006; Blüthgen *et al*., 2007; Dáttilo, Guimarães, & Izzo, 2013; Cagnolo & Tavella, 2015), mutualistic aphid-ant networks (Ivens *et al*., 2018) or parasite-ant networks (Elizalde *et al*., 2018). A notable exception is the global meta-analysis on different types of ant-symbiont interactions by Glasier, Poore, & Eldridge (2018). Unfortunately, this study only included a limited set of interaction types and pooled interactions of well-studied bioregions with those of very poorly studied regions.

Here, we ask what factors explain the topology of ant-symbiont networks across Europe. We firstly provide a quantitative and systematic meta-analysis on the diversity of European ant-symbiont interactions. By adopting the symbiont perspective, we test the hypothesis that the type of symbiosis explains the number and identity of their host species (host specificity). More specifically, we expect that parasitic ants are more specific than the other types of symbionts. In addition, we predict that different groups of symbiotic arthropods have highly variable host specificity. Secondly, we follow a trait-based host perspective to identify the major drivers that promote the diversity of ant-symbiont interactions and facilitate symbiont sharing. We test the hypothesis that the number of symbionts with which an ant species interacts and the number of symbionts it shares with other ant hosts depends on ecological factors (6 factors, including colony size, nest type, distribution, habitat, degree of sympatry, worker size) and evolutionary drivers (2 factors: phylogeny, biogeography) associated with the host species.

## II. MATERIAL AND METHODS

### (1) Ant symbionts

Symbionts are species that engage in a close association with a host species on which they may have beneficial, neutral or adverse effects. We limited our analyses to Europe (excluding the Canary Islands and Madeira), as ant-symbiont interactions in other continents are extremely fragmentary studied and poorly understood. Moreover a myriad of unknown symbionts are awaiting discovery and description in these continents (Parmentier, in press). By contrast, a firm body of knowledge on the distribution and diversity of ant symbionts in Europe has been recorded and has steadily grown from a long tradition of studying ant symbionts since the end of the 19th century (Wasmann, 1894; Janet, 1897). Depending on the intimacy of the relationship between ants and symbionts, we can distinguish obligate and facultative interactions. An obligate interaction occurs when a symbiont permanently lives inside or near an ant nest. Obligate symbionts completely depend on ants and cannot be found without them. Facultative myrmecophiles may associate with ants, but regularly (or mostly) occur without ants. In this study, we only focused on obligate symbionts.

We categorized five types of symbionts: (1) **Trophobionts** include aphids and scale insects that provide sugary honeydew in exchange of protection and hygienic services. These mutualistic arthropods are mostly found extranidally, except for aphids living on the roots of plants inside ant nests. (2) **Myrmecophiles ss** are a diverse group of arthropods that mostly live inside ant nests. Representatives of this group are distributed across many arthropod orders, but beetles and mites are the most diverse (Hölldobler & Wilson, 1990; Kronauer & Pierce, 2011). The life strategies of these organisms range from commensalism to specialized parasitism. (3) **Social parasites** encompass a group of ants that parasitize other ant species. There exist various types of social parasitism (see overview in Buschinger, 2009). In xenobiosis, cleptoparasitic ants construct a nest inside other ant nests, but raise their own brood. In temporary parasitism, a parasitic queen usurps a host colony and exploits the host work force to establish her own colony. Parasite workers will gradually substitute the host worker force. In dulosis (permanent parasitism with slavery), a parasite colony is established as in temporary parasitism. But here, the workers of the parasitic species will raid pupae of other ant species. Workers which will emerge from these pupae will do most of the tasks in the colony. In inquilinism (permanent parasitism without slavery), parasitic queens permanently exploit a host colony and no worker force is produced. The parasitic queen invests all her energy in producing sexuals. (4) ectoparasitic **Fungi**, (5) endoparasitic **Nematoda**. Plants engaging in mutualistic relationships (e.g. myrmecochory) were not included in our analyses. Contrary to the tropics, ant-plant relationships tend to be loose in Europe and are at best facultative.

### (2) Ant-symbiont dataset compilation

We compiled all documented ant-symbiont interactions in Europe. The database of ant-symbiont interactions was assembled from 253 published references, including faunistic notes, research articles, reviews and books. In a first round of searches, we scanned reference works (e.g. Wasmann, 1894; Donisthorpe, 1927; Evans & Till, 1966; Uppstrom, 2010; Tykarski, 2017; Molero-Baltanás *et al*. 2017) for associations between ant hosts and symbionts. Next we searched for ant-symbionts interactions via Google Scholar™ using the terms: “myrmecophile” or “ant associate” or “inquiline” or “ant guest” or “ant symbiont”. We also found host-symbiont interactions within references that were cited in the retrieved publications. In a second phase, each symbiont occurring in Europe was then searched by its Latin binomial name and its common taxonomic synonyms combined with a search string with the names of all ant genera (*N = 56*, AntWiki 2019) found in Europe (for example “Maculinea alcon” AND Acropyga OR Anochetus OR Aphaenogaster OR Bothriomyrmex OR Camponotus OR …) using Google Scholar. We chose Google Scholar over ISI Web of Science, as the latter does not retrieve faunistic notes or other types of grey literature. We omitted symbionts from our dataset when it was doubtful whether they are obligately associated with ants or not. Ultimately, we obtained a binary host-symbiont matrix filled with interactions (1s) and non-interactions (0s) between ants (columns) and symbionts (rows). Note that we included some references on ant-trophobiont interactions reported in the non-European part of Russia (e.g. Novgorodova, 2005) to ramp up the relative modest number of reported interactions in this type of association. The reported ants and trophobionts in these references have a widespread Palearctic distribution and they are expected to interact in Europe as well.

### (3) Host specificity in different symbiont types

We compared the number of host species across different types of host symbionts. Symbionts with only hosts identified at the genus level were not included in all following analyses. Symbionts are unevenly studied, which may result in better studied symbionts having a higher number of observed host species. To account for differences in sampling effort, we therefore first regressed the total number of host species against the (ln+1)-transformed number of google scholar hits for the binomial species name (and commonly used synonyms) of the symbionts. The residuals of this regression were then compared across symbiont groups using a GLM, followed by Tukey post-hoc tests in R 3.5.2.

In a second analysis, we wanted to test whether the relatedness of targeted host species varies among different symbiont groups. For each symbiont, we estimated the average taxonomic distance between the different hosts by using the specificity index, *S*_TD_ proposed by Poulin & Mouillot (2003). Host ant species (all ants belong to the family Formicidae) were classified following the Linnaean classification in subfamilies, tribes, genera and species groups/subgenera. The taxonomic distance between two hosts is then defined as the number of hierarchical steps that are needed to reach a common node in the taxonomic tree. The taxonomic distance between two species of the same subgenus/species group equals to 1, the distance between two species of the same genus, but from a different subgenus/species group equals to 2. A maximum taxonomic distance of 5 is reached between two ant host species from different subfamilies. *S*_TD_ was estimated by averaging the taxonomic distance across all pairs of host species. We modelled the specificity index against symbiont type with a GLM followed by Tukey post-hoc tests. Symbiont species with a single reported host species were omitted from this analysis, as no specificity index can be estimated for these species (a total of 382 symbionts were retained in this analysis). In parallel, we reran this analysis with a phylogenetic instead of a taxonomic distance matrix. The phylogenetic distance matrix was based on the phylogenetic tree of European ants by Arnan *et al*. (2017). Distances between species were estimated by node count (number of nodes along the branches of the tree from one species to another) and were retrieved using Mesquite v.3.5. Phylogenetic distances are more accurate than taxonomic distances to assess relatedness, but unfortunately we do not possess phylogenetic information at the species level for all ants in our dataset (phylogeny of 96 out of 177 ant species was known). We decided to drop the 81 ant species without phylogenetic information and the corresponding interactions with their symbionts for subsequent analyses. We believe that this is acceptable as these species only cover 13.5% of the interactions in our host-symbiont dataset. We also omitted the exotic ant species *Linepithema humile*, consequently we reduced our dataset to 95 ant species and 602 symbionts. In addition, symbionts which interacted with only one ant species were omitted, as no specificity index can be calculated for these species. Ultimately, we retained 349 symbiont species in this analysis.

### (4) Predictors of symbiont diversity in European ants

A central question in this study is why some ant species host more symbionts than other ant species. Therefore, we first compiled for the European ant species several predictors, based on Arnan *et al*. (2017), Boulay *et al*. (2017), Seifert (2007) and AntWiki (2019) reflecting differences in host functional niche. Predictors of symbiont diversity were extended traits of the ant species and colony, including colony size (number of workers), average worker size (mm), nest type (levels: arboricolous, diverse, soil and organic mound) and phylogeny as a proxy for trait similarity, and ecological factors of the ant species, including habitat (levels: eurytope, introduced exote, open, open/sylvicolous and sylvicolous), distribution range, degree of sympatry and biogeographic region (levels: Atlantic, Boreomountain, Continental, Mediterranean, wide-ranging). Ants were assigned to the biogeographical region where they were found proportionally the most in sampled biogeographical regions based on the observations of Arnan *et al*. (2017). If the proportional occurrence in the most preferred region was less than double of the proportional occurrence in another region, the ant species was grouped under the “wide-ranging” category. We also estimated the distribution range (the number of countries where the host species has been reported, based on records on AntWiki, 2019), the degree of sympatry (number of ant species with symbionts which share at least one country in the distribution range, based on AntWiki, 2019) and the number of hits for their binomial name (and common synonyms) on google scholar as a proxy for sample effort for every ant species. Next we correlated total symbiont diversity with the host predictors described above, while correcting for the phylogenetic relatedness of the different ant species. The phylogenetic relatedness of host species should be accounted as closely related host species cannot be treated as independent observations. For that reason, we modelled a phylogenetic generalized least squares regression (PGLS) with total number of symbionts per ant species as dependent variable. A PGLS model incorporates a phylogenetic variance-covariance matrix in its error structure. We used the variance-covariance matrix based on the pairwise node counts retrieved from the phylogenetic tree of European ants by Arnan *et al*. (2017). The phylogenetic relatedness of 96 out of the 177 ant species found in our dataset was determined in this tree. We decided to drop the 81 ant species without phylogenetic information and the corresponding interactions with their symbionts for subsequent analyses. We believe that this is acceptable as these species only cover 13.5% of the interactions in our host-symbiont dataset. We also omitted the exotic ant species *Linepithema humile*, consequently we retained a dataset with 95 ant species and 602 symbionts. The phylogenetic covariance matrix was multiplied by Pagel’s λ, a widely used parameter that scales and corrects for the expected degree of phylogenetic covariance (Pagel, 1999). This multiplier spans from 0, which corresponds with the complete absence of phylogenetic signal in the residuals of the model (the model is then similar to a regular GLM with an ordinary least squares error structure) to 1, when the covariance of the model residuals follows a Brownian motion model of evolution (Pagel, 1999; Freckleton, Harvey, & Pagel, 2002). The λ parameter characterizing the phylogenetic signal was estimated through maximum likelihood estimation within the PGLS model. We analysed this model using the *pgls* function embedded in the R 3.5.1-package ‘caper’. We transformed the variables to meet the normality assumptions of the residuals. Number of symbionts was fourth root transformed, the predictors colony size, degree of sympatry and google scholar hits ln-transformed, and the distribution range was square root transformed. Finally, all continuous predictors were scaled to unit variance.

In addition to this analysis on the drivers of total symbiont diversity, we ran similar PGLS models with subsets of symbiont species richness as dependent variables (overall number of myrmecophiles, trophobionts, social parasites, and the diversity of the largest groups of myrmecophiles, i.e. beetles and mites, separately) and predictors of the subset of ant species that engage with these symbionts as predictors. Identical transformations of predictors and subsets of symbiont richness were applied as in the analysis on total symbiont richness described above. Diversity of social parasites and fungi were not regressed against ant predictors in separate PGLS models as the number of host ants are relatively low in these groups.

Models were ranked per analysis with the *dredge* function in ‘MuMIn’ R-package according to their AICc-value (corrected Akaike Information Criterion). We retained the best models identified with Δ AICc < 2. Significance levels of the predictors of the retained models were assessed using Likelihood ratio tests.

### (5) Predictors of symbiont sharing in European ants

Understanding the factors that facilitate or constrain the transmission of a symbiont from one host to another, is pivotal in the study of host-symbiont networks. Here we assessed which drivers promote the sharing of symbionts in ants. Predictors were similar to the previous analysis and encompassed extended traits of the ant species and colony, including colony size, worker size, nest type (levels: arboricolous, diverse, soil and organic mound) and phylogeny as a proxy for trait similarity; and ecological factors of the ant species, including habitat (levels: eurytope, introduced exote, open, open/sylvicolous and sylvicolous), distribution range, biogeographic region (levels: Atlantic, Boreomountain, Continental, Mediterranean, wide-ranging) and degree of sympatry. We used multiple regression on distance matrices (MRM), an extension of partial Mantel analysis, to test the association between different distance matrices (Lichstein, 2007). The dissimilarity symbiont matrix contained the pairwise Jaccard distances between each pair of host ants based on the presence-absence data of the symbionts they supported. This matrix was regressed against multiple distance matrices giving dissimilarities in the aforementioned predictors. Worker size distance was the absolute difference for this trait between every pair of ant species. The pairwise differences in colony size were ln-transformed. The sympatry matrix listed the ln-transformed number of countries in the distribution range that are shared between each pair of ant species. Entries in the distance matrices of habitat, nest type and biogeographic region were coded 0 when the pair of ants occupy the same habitat, nest type or biogeographic region, respectively, and 1 when the pair of ants show differences in these variables. The phylogenetic distances were the pairwise node counts. Again, we focused our analysis on the subset of 96 ants for which the phylogenetic relationship was resolved by Arnan *et al*. (2017). We also included a matrix of sampling effort in which we pairwise multiplied the (ln+1)-transformed hits on google scholar of one ant species with the (ln+1)-transformed hits on google scholar of another ant species. All matrices were standardized between 0 and 1 and Multiple Regression On Distance Matrices (MRM)-analyses were conducted in the R package ‘ecodist’ using the MRM function. Significance of the predictor matrices was tested using 9999 permutations. We removed non-significant predictors, and reran the MRM analysis until all predictors were significant (Martiny *et al*., 2011). The relative importance of the significant predictor matrices was calculated with the lmg-metric, which uses unweighted averages of sequential R^2^ of different orderings of the model predictors. The calculation and visualization of the lmg-metrics was conducted with the R-package ‘relaimpo’.

We conducted similar MRM-analyses on subsets of the symbiont community, where the response variable was the dissimilarity (pairwise Jaccard indices) in the set of myrmecophiles, trophobionts, social parasites, myrmecophilous beetles and myrmecophilous mites of the host ants, respectively.

## III. RESULTS

### (1) Ant-symbiont networks display a diversity of species interactions

We identified 701 obligate ant symbionts interacting with 177 ant species in Europe (Appendix S1). The references we used to compile the host-symbiont interaction matrix are listed per symbiont species in Appendix S2. Myrmecophiles (*N* = 537) outnumbered the four other groups of ant symbionts (*N* = 71, *N =* 69, *N =* 13 and *N* = 11 for respectively social parasites, trophobionts, ectoparasites (all fungi) and endoparasites (all nematodes in our analysis). Within the group of myrmecophiles, beetles and mites were the most species-rich groups (Fig. 1). The hosts of 73 symbionts were not identified at the species level in the literature record. The distribution of the number of host species per symbiont was right-skewed (mean = 3.58, median = 2). The highest frequency (39%) of symbionts interacted with 1 host species and a maximum number of 34 host species was documented in the myrmecophilous silverfish *Proatelurina pseudolepisma*.

**FIGURE 1.**
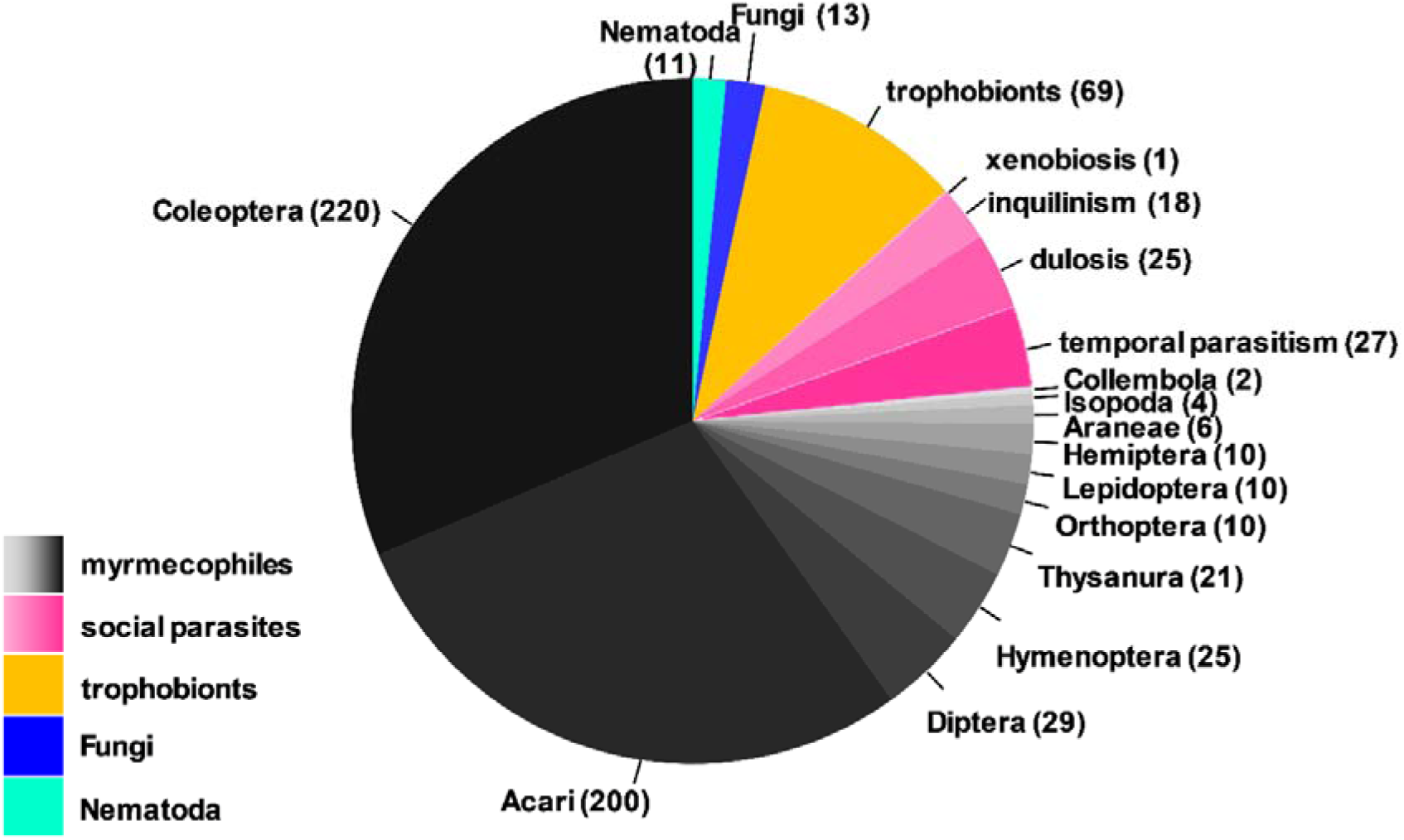
Distribution of different types of ant symbionts. Total number of symbionts *N =* 701, number of symbionts per type given in brackets.

European ant genera contained a highly variable number of species (proportional to the genus font size in Fig. 2, Fig. S1), ranging from 1 to 82 species, in the genera that interact with symbionts. Generally, the species-rich ant genera, such as the Formicinae genera *Lasius*, *Formica* and *Camponotus* and the *Myrmicinae* genera *Messor* and *Myrmica* attracted a higher diversity of all five types of symbionts (Pearson’s product-moment correlation: r = 0.60, P < 0.001, df = 28, Fig. S1). A notable exception was the species-richest European ant genus *Temnothorax*, which supported a relatively moderate number of symbionts. Myrmecophiles were the dominant group in most ant genera (mean 63.7% of the total symbiont diversity). Trophobionts were generally the second most diverse group (mean 14.9%), but were absent or nearly absent in some genera such as *Monomorium*, *Aphaenogaster*, *Leptothorax, Messor* and *Cataglyphis*. Social parasites slightly contributed to total symbiont diversity in most ant genera (mean 8.9%), but were very diverse in the ant genera *Temnothorax*, *Tertramorium* and *Leptothorax*. Fungi (mean: 11.7%) and nematodes (mean: 0.9%) are also small symbiont groups. Interestingly, *Myrmica* is targeted by a relatively high number of ectoparasitic fungi. Ant genera shared many symbionts (74.2% on average) with other genera, belonging to the same or different ant subfamilies (connecting lines in Fig. 2). *Temnothorax*, *Leptothorax* and *Messor* are characterized by a relatively high number of unique symbionts (cf. relative large inner circles in Fig. 2).

**FIGURE 2.**
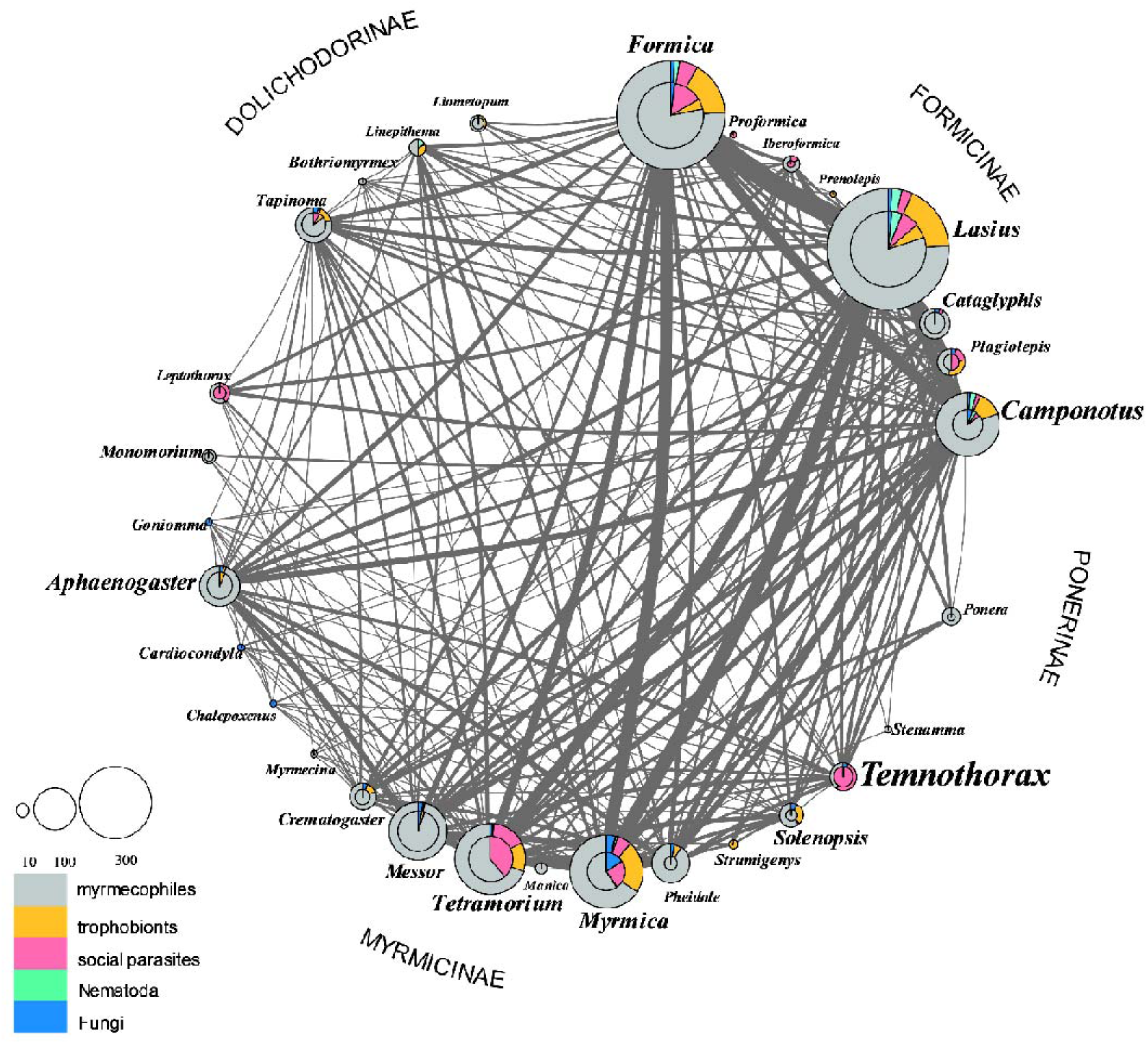
Ant symbiont network displaying the proportional distribution of symbionts across the European ant genera. A multilevel pie chart is given for each ant genus. The size of the outer pie chart corresponds to the total number of associated symbionts. The size of the inner pie chart is related to the number of symbionts that are not shared with other ant genera (unique symbionts). The proportional distribution of the five types of ant symbionts is given for all associated symbionts (colour segments in outer pie charts) and for the symbionts that are not shared with other genera (colour slices in inner pie charts). The relative proportion of unique symbionts can be deduced by the relative size of the inner circle to the outer circle. The genera are organized in four groups, corresponding to the ant subfamily to which they belong. The genera are connected with lines, of which the width is directly proportional to the number of shared symbionts. The font size of a genus is proportional to its number of described species in Europe.

### (2) Host specificity in different symbiont types

After controlling for sampling effort, host range of the symbiont groups was significantly different (GLM, df = 25, P < 0.001, Fig. 3). Social parasites targeted the lowest number of host species, whereas myrmecophilous Thysanura, Collembola and Isopoda are associated with the highest number of ant hosts. Trophobionts also interacted with a relatively high number of ant hosts (Fig. 3). Host range of symbiont groups without controlling for sampling effort can be found in Fig. S2.

**FIGURE 3.**
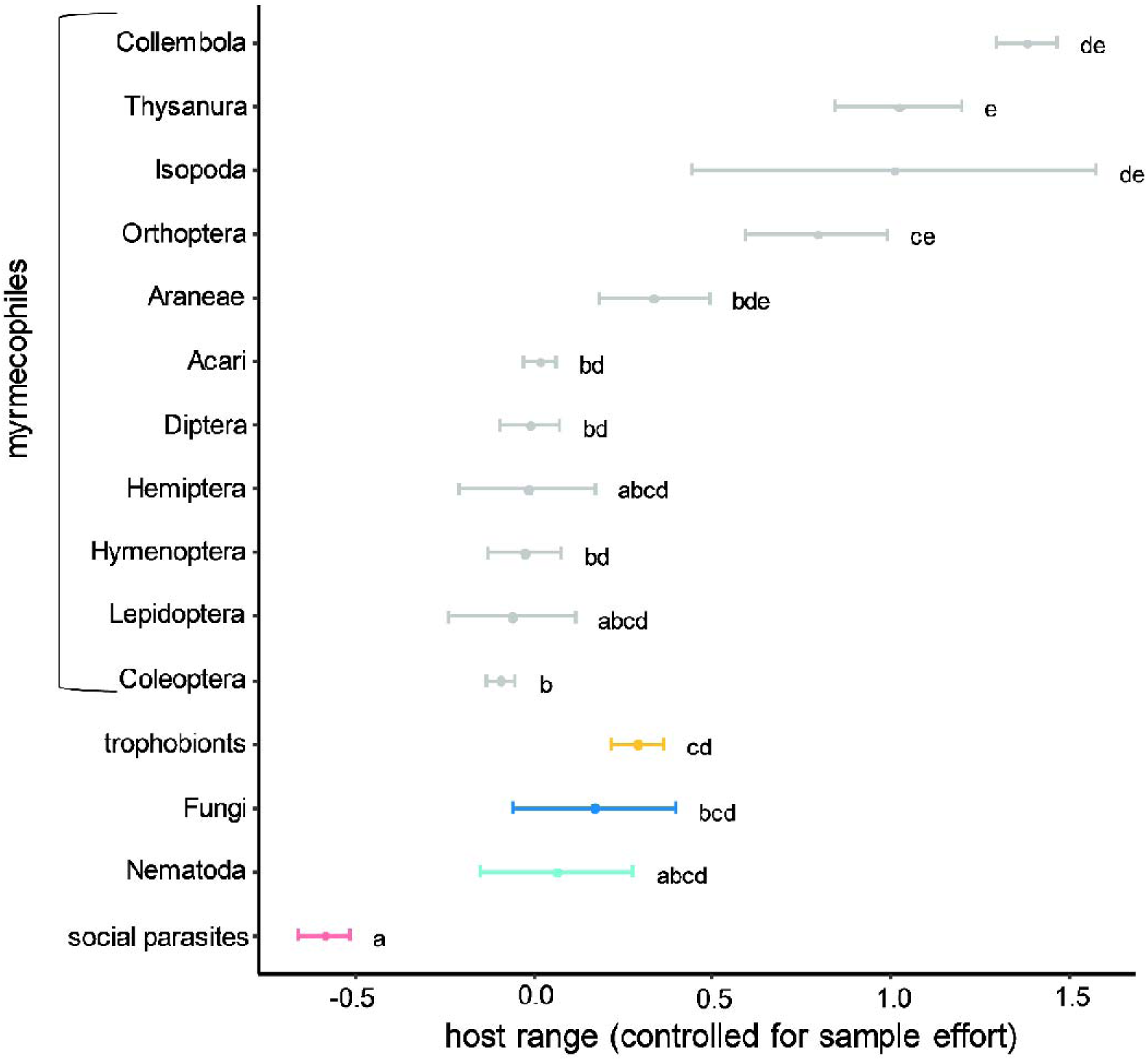
Number of host species per symbiont type, controlled for sampling effort ± SE. The different myrmecophilous arthropod orders are displayed in grey. Letter codes refer to Tukey post-hoc test. Species with no letters in common are significant different P < 0.05.

The average taxonomic distance between host species targeted by a symbiont was significantly different among the symbiont groups (GLM, F = 9.4, df = 14, P *<* 0.001, Fig. 4). Likewise, we found significant differences in average phylogenetic distance between host species across the symbiont groups (GLM, F = 9.4, df = 14, P *<* 0.001, Fig. S3). Patterns in the average host phylogenetic distance of the different symbiont groups were closely matching those of the average host taxonomic distance. The host species of social parasites were very closely related to each other (Fig. 4 and S3). Fungi also exploited related host species. Trophobionts, ectoparasitic nematodes and myrmecophiles interacted with hosts that are much more unrelated. The group of myrmecophiles was heterogeneous and when we subdivided it into different subgroups, we obtained an entire gradient in average host taxonomic/phylogenetic distance (Fig. 4 and S3).

**FIGURE 4.**
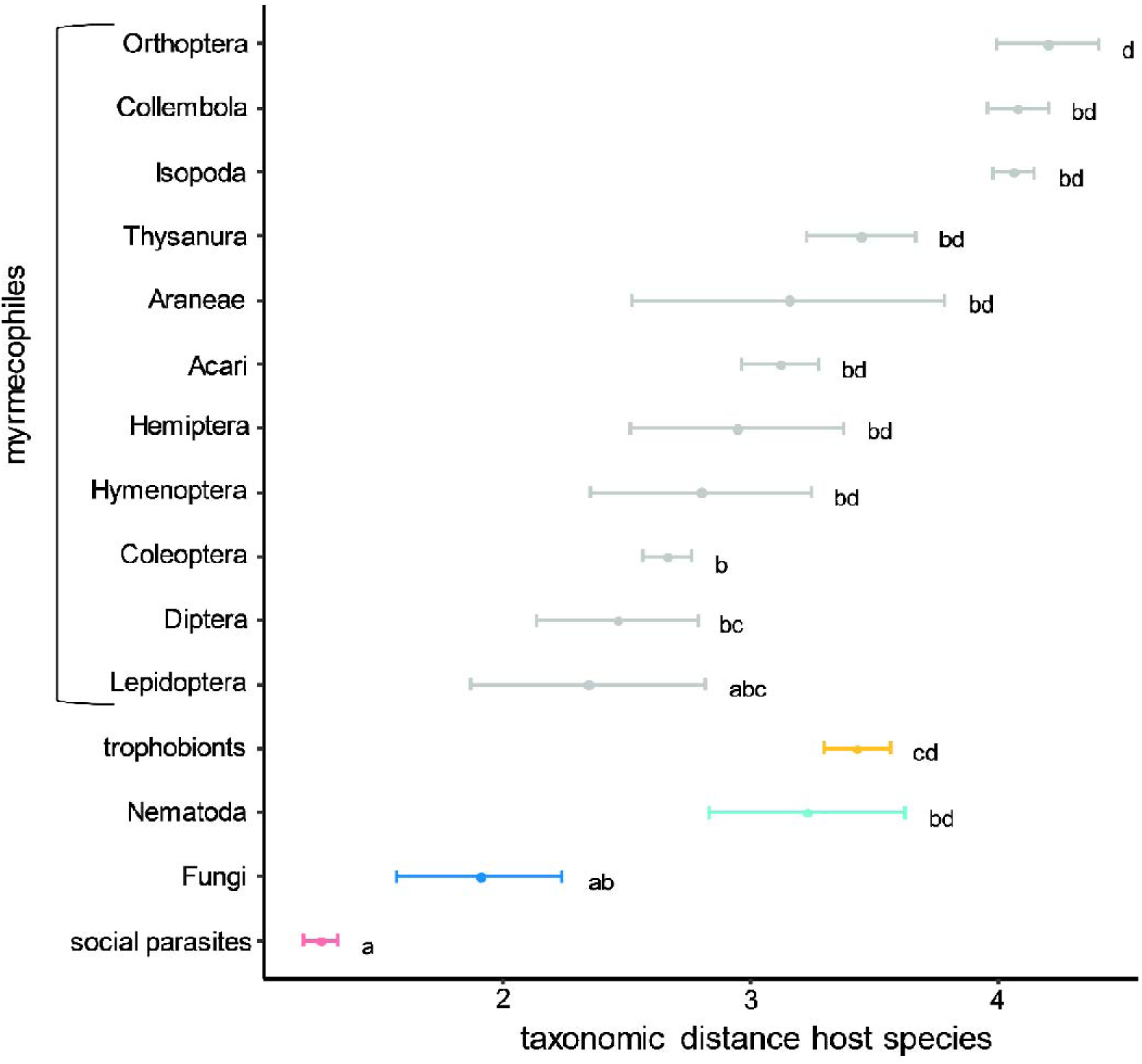
Taxonomic distance ± SE of targeted host species for different types of ant symbionts. The different myrmecophilous arthropod orders are displayed in grey. Letter codes refer to Tukey post-hoc test. Species with no letters in common are significant different P < 0.05.

### (3) Predictors of symbiont diversity in European ants

The number of symbionts is highly variable in ant species. Here we report the host drivers that affect total symbiont richness and diversity of subsets of ant symbionts (myrmecophiles, trophobionts, social parasites, myrmecophilous mites and myrmecophilous beetles). Total symbiont diversity was clearly positively correlated with colony size. This factor was highly significant (PGLS, P < 0.001) in the 4 top-ranking models (Table 1, Fig. 5). Habitat and distribution range of the host were also incorporated in most of the top-ranking models. In these models, symbiont richness increased with the host distribution range and was highest in eurytopic habitats (PGLS, P < 0.001). As expected, sample effort has a major effect on the reported total symbiont diversity and the five subsets of symbiont diversity. Symbiont interactions were the highest in ants that are intensively studied. We controlled for sample effort by including the proxy (ln+1)-transformed google scholar hits in our models. Myrmecophile richness was also positively affected by colony size (PGLS, P < 0.001 in the 5 top-ranking models, Table 1, Fig. 5), distribution and eurytopic habitat. The diversity of myrmecophilous beetles was positively correlated with colony size and distribution range of the host (PGLS, P-values < 0.001). Myrmecophilous mite diversity was also positively correlated with colony size and distribution range (PGLS, P-values < 0.001, Table 1) in all retained models. Trophobiont diversity was mainly driven by sampling effort (PGLS, P < 0.001, Fig. 5, Table 1). Trophobiont diversity was consistently highest in the Atlantic region and in ant species with small workers. There were no predictors consistently present in the top-ranking models explaining social parasite species richness (Table 1).

**FIGURE 5.**
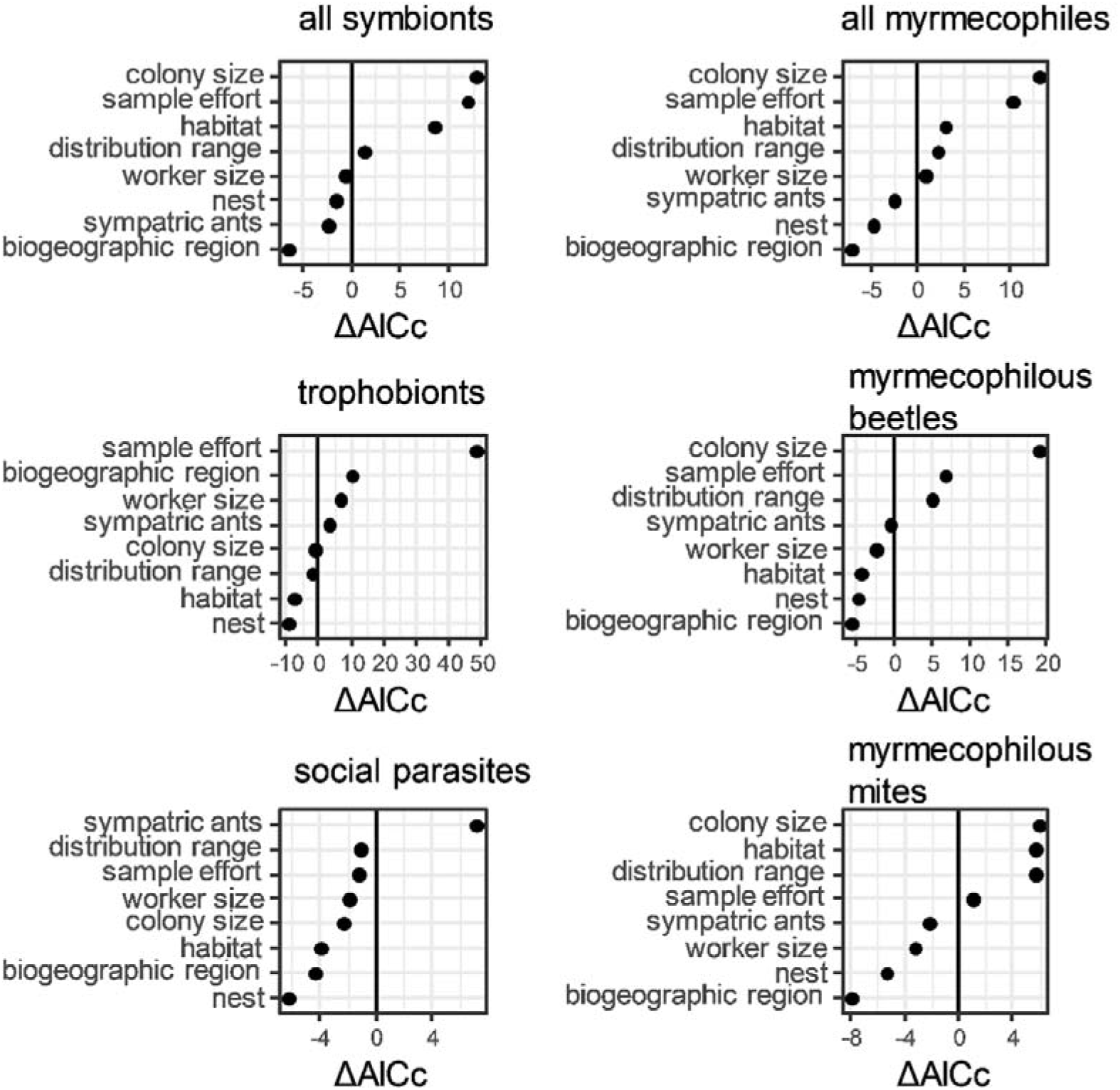
Ranking of the predictors from the six PGLS models by the corrected Akaike information criterion (ΔAICc). The change in AICc when adding or removing a variable from the most optimal model is compared. Predictors included in the most optimal model are removed (ΔAICc positive), whereas those not included are added (ΔAICc negative) to the best model (lowest AICc). The ranking is given for the six PGLS analyses, i.e. with dependent variable the number of symbionts (best model: ∼ sample effort + colony size + distribution + habitat), trophobionts (best model: ∼ sample effort + biogeographic region + worker size + sympatric ants), social parasites (best model: ∼sympatric ants), myrmecophiles (best model: ∼ sample effort + colony size + distribution + habitat), myrmecophilous beetles (Coleoptera) (best model: ∼ colony size + sample effort + distribution range) and myrmecophilous mites (Acari) (best model: ∼ colony size + habitat + distribution range + sample effort), respectively. Note that mites and beetles are nested within the group of myrmecophiles and myrmecophiles, trophobionts and social parasites are three subsets of all ant-associated symbionts.

**Table 1.**
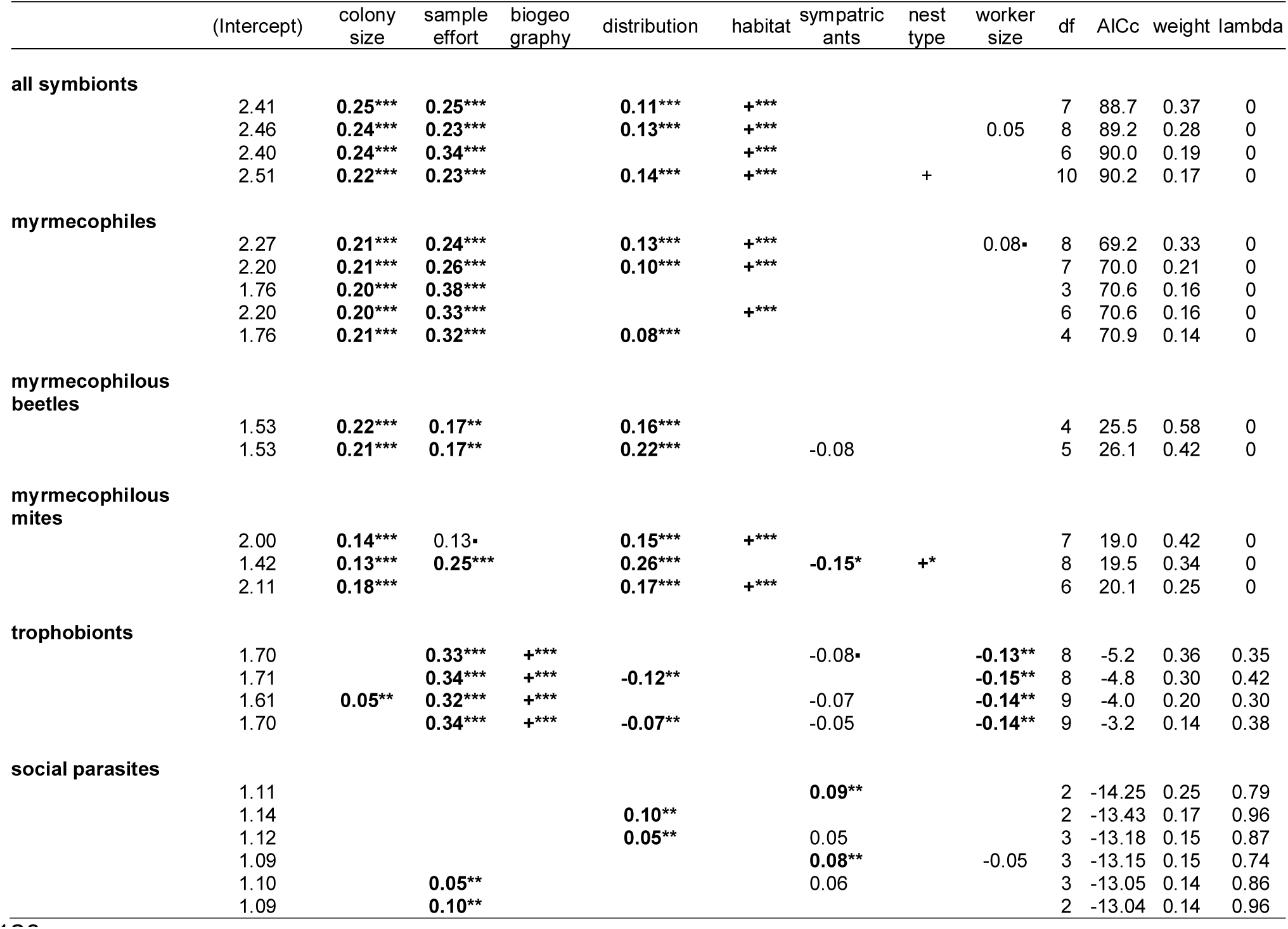
Estimates for the predictors of the top-ranked PGLS-analyses (ΔAICc < 2). The subset of best models is given for the analyses with total number of (1) symbionts, (2) myrmecophiles, (3) myrmecophilous beetles, (4) myrmecophilous mites, (5) trophobionts and (6) social parasites as dependent variable. Significant estimates indicated in bold (P < 0.001: ***, P < 0.01: **, P < 0.05: *, P < 0.10: ^▪^).

We did not find a phylogenetic signal in the predictors of the PGLS models (Δ AICc < 2) with total symbiont, total myrmecophile, myrmecophilous beetle and myrmecophilous mite diversity as dependent variables. The residuals of these models showed no phylogenetic covariance (Pagel’s λ lambda was 0, Table 1). In contrast, phylogenetic relatedness of the hosts explained additional variation in the social parasite models (Pagel’s λ ranged between 0.74 and 0.96) and to a lesser degree in the trophobiont models (Pagel λ = 0.30-0.42) (Table 1).

### (4) Predictors of symbiont sharing in European ants

Here we examined whether the sharing of symbionts is affected by the similarity in particular traits and ecological drivers of the host species. The maximum amount of variation explained by our predictor matrices was in the subset of trophobionts (R^2^ = 37.1%, Fig. 6). MRM analyses aiming to explain similarities in particular groups of symbionts explained more variation (R^2^ ranging from 14.7% to 37.1%) than those focusing on similarities in the total symbiont community (R^2^ = 9.8%). Sample effort positively affected similarities in symbiont communities including all symbionts (MRM, proportional contribution to the total R^2^: lmg = 0.11, P < 0.001). Well-studied ant pairs also shared more trophobionts, social parasites, myrmecophiles, myrmecophilous beetles and mites (MRM-analyses, lmg ranging from 0.18 - 0.58, all P-values < 0.01, Fig. 6). The most important predictors of similarity (dissimilarity) in ant symbiont communities in the European ant dataset were phylogenetic relatedness of the ant hosts (1-distance) (MRM, lmg = 0.40, P = 0.001) and similarity (1-dissimilarity) in biogeographic region (MRM, lmg = 0.35, P = 0.001) (Fig. 6). Similarities in worker size and colony size also facilitated the sharing of symbionts (MRM, lmg = 0.08, P = 0.01 and lmg = 0.05, P = 0.02, respectively, Fig. 6). Trophobiont sharing was also positively correlated with phylogenetic relatedness and similarity in biogeographic regions of the ant hosts (MRM, lmg = 0.26, P < 0.001 and lmg = 0.16, P < 0.001, resp.). The similarities in social parasite communities was largely explained by phylogenetic relatedness (MRM, lmg = 0.53, P < 0.001). Similarities in biogeography and colony size explained additional variation in the sharing of social parasites (MRM, lmg = 0.20, P < 0.001 and lmg = 0.05, P < 0.05). Interestingly, phylogenetic relatedness of the hosts did not promote the sharing of myrmecophiles and myrmecophilous mites. Host relatedness had only a minor effect on myrmecophilous beetle sharing (MRM, lmg = 0.02, P = 0.03). The assemblage of myrmecophiles, myrmecophilous beetles and mites was similar in pairs of ants occupying identical biogeographic regions (MRM, lmg = 0.57, P < 0.001 and lmg = 0.38, P < 0.001 and lmg = 0.16, P = 0.02, resp.) and having a large overlap in distribution (degree of sympatry) (MRM, lmg = 0.13, P = 0.04 and lmg = 0.24, P = 0.005 and lmg = 0.28, P = 0.002, resp.) (Fig. 6).

**FIGURE 6.**
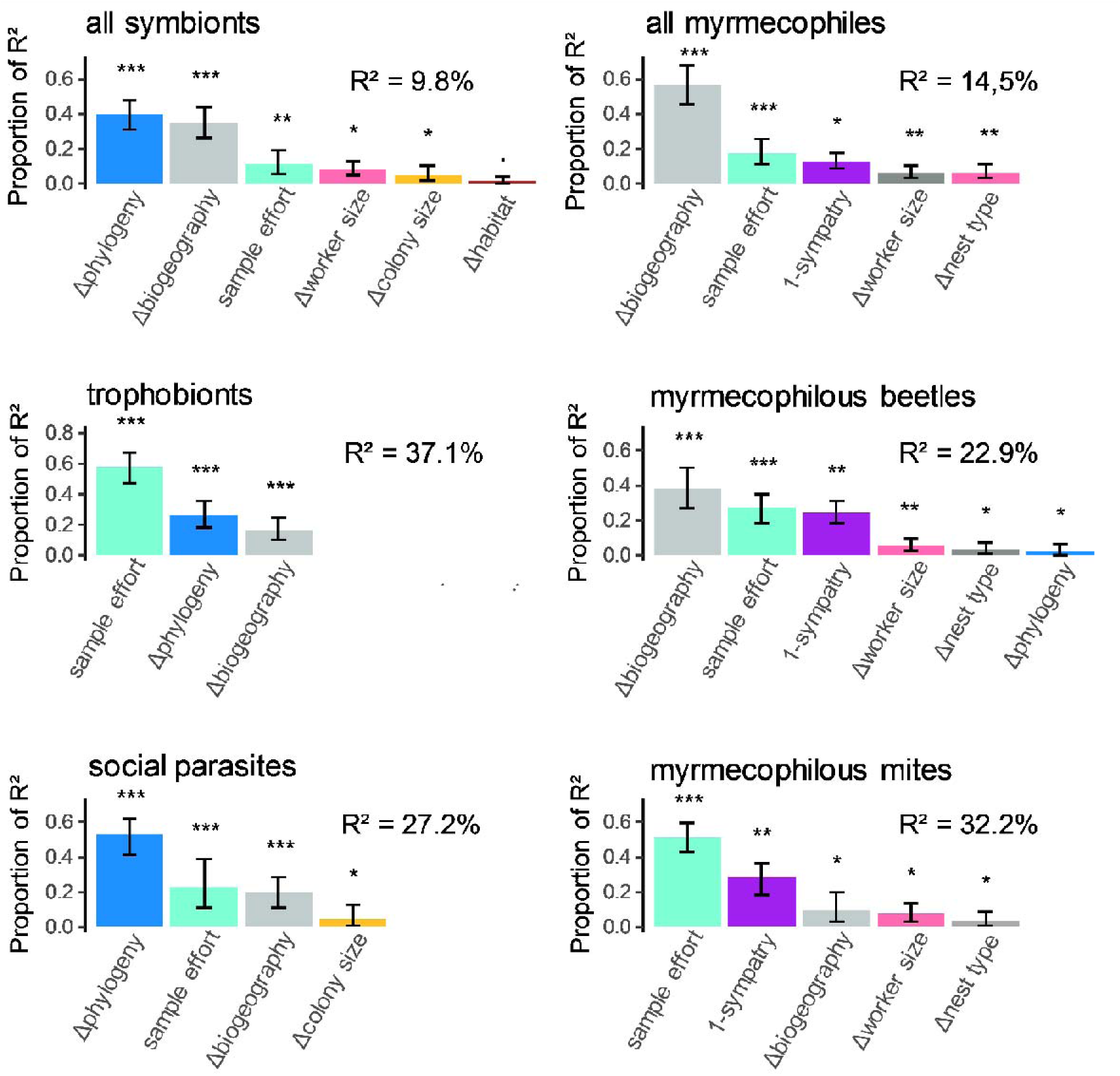
Relative importance of the significant predictor matrices explaining the dissimilarity in symbiont communities across different ant species. Rankings are given for predictors explaining overall dissimilarity (1-similarity) in symbiont composition, and for dissimilarity in subsets of symbiont composition: trophobionts, social parasites, myrmecophiles, myrmecophilous beetles and mites, respectively. Note that beetles and mites are nested within the group of myrmecophiles, and myrmecophiles, trophobionts and social parasites are three subsets of all ant-associated symbionts. The allocated contribution (sequential R^2^) of the different distance matrices (indicated with Δ) or other predictor matrices (sample effort and 1-sympatry) to the explained variation of the MRM models is estimated with the lmg-metric. All predictor matrices are positively correlated with dissimilarity in symbiont composition. Significance levels of the predictors were tested with a permutation test (n = 9999, P < 0.001: ***, P < 0.01: **, P < 0.05: *, P < 0.10: ^▪^).

## IV. DISCUSSION

Understanding the patterns of community assembly has been a central topic in ecology. Community structuring and the underlying species interactions have been increasingly been approached using network analysis. Yet, studies that describe and aim to understand the topology of large-scale ecological networks consisting of different interaction types are limited. We here provide a complete tally of the distribution of ant symbiont groups over European ants and compare host specificity, symbiont richness, host switching and its drivers for five different ant symbiont groups.

### (1) Characterization of the European ant-symbiont network

It is widely acknowledged that the group of obligate ant symbionts is hyperdiverse (Wasmann, 1894; Kistner, 1979, 1982; Hölldobler & Wilson, 1990; Rettenmeyer *et al*., 2010), although exact species numbers at a regional scale are lacking. Rough estimates of the global diversity of parasites living in ant nests reach 10.000 to 20.000 species (Thomas, Schönrogge, & Elmes, 2005), which is higher than mammal and bird diversity. We here identified 701 obligate symbionts distributed over 177 ant species in Europe. The majority of the symbionts were classified as myrmecophiles, which are commensalistic to parasitic arthropods mostly living inside the nest of ants (Kronauer & Pierce, 2011; Parmentier *et al*., 2016; Parmentier, in press). The large diversity of this group is fuelled by the species-rich myrmecophilous beetle and mite communities. In other regions, beetles and mites outnumber other arthropod groups as well (Kistner, 1982; Hölldobler & Wilson, 1990). Social parasites and mutualistic trophobionts are medium-sized groups; parasitic nematodes and fungi are relatively species-poor, but definitely understudied. Mutualistic ant symbionts are thus clearly overshadowed by the diversity of commensalistic and parasitic ant symbionts in Europe. Speciose ant genera and subfamilies generally supported higher numbers of ant symbionts. Host-symbiont networks are characterized by an asymmetrical organization of interactions with host-specific symbionts and symbionts that interact with multiple host taxa (Guimaraes *et al*., 2006). Overall, a large proportion of the symbionts were shared among heterogeneric ant species (Fig. 2). Some ant genera interacted with a relatively low number of symbionts, but most of their symbionts were not found in association with other ant genera. The highly specific group of social parasites was much more represented in the symbiont community of these hosts. In addition, the group of trophobionts is diverse in some ant genera, but it is marginal or even absent in others (Fig. 2). The uneven distribution of the five types of symbionts among the ant genera suggest that some ant lineages are more predisposed to associate with particular types of symbionts. Ant-host associations are thus shaped by deep evolutionary processes as determined by biotic and environmental drivers of speciation and extinction (Aguilée *et al*., 2018).

### (2) Host specificity in different symbiont types

Host specificity is a key feature of host-symbiont networks, and is moulded by the ecological and evolutionary interactions between host and symbiont (Poulin & Mouillot, 2003). Patterns in host specificity have been studied in a wide range of host-symbiont systems. Generally, parasites are thought to have a tendency to evolve to extreme host specialization as they need complex adaptations to by-pass host defence (Kawakita *et al*., 2010). The drivers favouring host specificity in mutualist partners are far less understood and both low and high degrees of specificity are widespread (e.g. plant-seed dispersers and fig-fig wasps, respectively). Ant symbioses are ideal to unravel patterns in host specificity. They occupy the complete mutualism-parasitism continuum and allow comparison of host specificity in different types of symbionts. We here demonstrate that average host range in European ant symbionts was much broader than previously assessed in a study on host specificity of myrmecophiles at a global scale (Glasier *et al*., 2018). The authors of this study state that obligate ant symbionts occurred on average with ca. 1.20 host species. We, however, found that European symbionts were reported with three times this number of host species (3.57) on average. The extreme low number of the detected hosts in the study of Glasier *et al*. (2018) is the result of their searching method. They overlooked data hidden in faunistic notes, grey literature and books, which report the majority of the interactions between ants and their symbionts. Moreover, the symbiont fauna, let alone the range of their interactions, is poorly documented outside Europe, which makes hard predictions unreliable (Parmentier, in press). Ant symbionts were extremely variable in ant host range. After controlling for sample effort, social parasites clearly targeted the lowest number of host species, which is in line with the expectations as they are the most specialized group of parasites (Buschinger, 2009). The commensalistic myrmecophilous group of silverfish and the mutualistic trophobionts engaged symbioses with the highest number of ant species. Apart from the quantity of the hosts, the relatedness of the host species is a vital aspect of host specificity. It is well described that social parasites colonize nests of related hosts (Buschinger, 2009). However, this has never been compared with other type of symbiont groups. We showed that the hosts of social parasites were clearly the most related of all symbiont types. The hosts of myrmecophiles groups showed moderate to poor relatedness on average. Trophobionts and nematodes associated with distantly related ant species.

There is a large body of literature that explains the constraints of host switching in social insect symbionts. Generally, it is thought that host-specific myrmecophiles and social parasites rely on chemical deception, by mimicking the colony recognition cues or some key pheromones (overview in Parmentier, Dekoninck, & Wenseleers, 2017). They are completely integrated in the host colony and are treated as a true colony member. Because of the strict mimicking of the host’s unique chemical profile, they are not able to colonize unrelated host species. Generalist myrmecophiles are typically poorly integrated in the colony, but host switching is more common in this group. It is facilitated by the use of general defensive chemicals, chemical insignificance or behavioural strategies (Stoeffler, Tolasch, & Steidle, 2011; Parmentier *et al*. 2017, 2018). The processes which make aphids attractive for one host, but not for another are hitherto unexplored. Likewise, the factors that facilitate or constrain host switching in parasitic nematodes and fungi are unknown.

### (3) Predictors of symbiont diversity in European ants

Associations between ant hosts and their symbionts are not random and structured according to both ecological and evolutionary factors that act at different spatiotemporal scales. From the perspective of ant symbionts, ant nests can be conceptualized as islands or local patches connected through dispersal. In analogy with McArthur & Wilson’s island theory and Leibold’s metacommunity framework, larger ant nests interact with more symbiont species. Ant nest size was repeatedly hypothesized as an important driver of ant symbiont diversity (Hughes, Pierce, & Boomsma, 2008; Kronauer & Pierce, 2011), but here formally tested for the first time. Previous studies across very different host-symbiont systems (e.g. ectoparasites of fishes in Guégan *et al*., 1992, parasites of hoofed mammals in Ezenwa *et al*., 2006, parasites of carnivores Lindenfors *et al*., 2007, feather mites of finches Villa *et al*., 2013) identified host body size as one of the key factors in determining symbiont species richness (Kamiya *et al*., 2014). This positive association results from the fact that larger hosts provide more niches and are less ephemeral (Lindenfors *et al*., 2007). Analogously, larger ant nests provide more space to allow larger population sizes and thereby reducing the extinction risk of symbionts. In addition, larger nests contain a higher diversity of microhabitats, including refuge areas that eventually facilitate species coexistence (Barabás, D’Andrea, & Stump, 2018). Larger ant nests are also expected to be more persistent (Kaspari & Vargo, 1995). Colony size is thus a strong local driver of symbiont richness and holding for myrmecophiles in particular. Myrmecophile diversity is additionally strongly determined by more regional ecological factors like range size and niche width of the host ants, so factors related to their general abundance and distribution. In that respect, eurytopic ants, such as *Lasius niger* and *Myrmica rubra* that can live in a wide variety of habitats including urban regions, hosted more symbionts. Both the effect of distribution and habitat reflect that more symbionts occur in widely distributed ant species with high densities. Symbionts associated with widely distributed ants are less prone to extinction as predicted by life history theory and metacommunity ecology (Nosil, 2002; Leibold *et al*., 2004). Host density has widely been demonstrated as another key factor for explaining parasite species richness (Lindenfors *et al*., 2007). Together with local colony sizes, sampling effects appear highly relevant, hence more neutral drivers, of their diversity among different hosts. Interestingly, trophobiotic and more specialised parasitic interactions as seen in the group of social parasites are more determined by evolutionary drivers. Indeed, affinity with their hosts is not only shaped by biogeography, hence a common spatial evolutionary ancestry but also by phylogeny. The effect of phylogeny is echoed in the high Pagel’s lambda values of the corresponding PGLS models, implying that much of the residual variation in trophobiont and especially social parasite richness could be explained by the phylogeny of the hosts. The strong phylogenetic driver for social parasite richness is in line with our previous results that social parasites mainly target closely related ant species (referred to as Emery’s rule, Buschinger, 2009) belonging to a small number of ant genera. Symbiont network structure thus shifts from more neutral ecological drivers related to regional species abundance to co-evolutionary drivers related to ancestry. The uniqueness and tightness of species interaction are known to be both a driver and consequence of co-evolutionary dynamics. Interestingly, we here show these evolutionary drivers to overrule any ecological one in the most antagonistic interactions, hence demonstrating the integrated nature of symbiont network formation according to the prevailing interaction strengths. Ant-symbiont networks are quite unique in the sense that the host associated network that is studied is covering a wide array of interactions, from putatively mutualists to strictly antagonistic. Studies on specialised host-parasite networks equally point at the dominance of evolutionary drivers (phylogeny and biogeography) of these associations (Feliu *et al*., 1997; Rosas-Valdez & de León, 2011), while less obligatory animal parasitic (Nunn *et al*., 2003; Ezenwa *et al*., 2006; Lindenfors *et al*., 2007; Nava & Guglielmone, 2013) or plant mutualistic interactions (Sanders, 2003; Wagner, Mendieta-Leiva, & Zotz, 2015) show more ecological factors related to distribution and abundance patterns that enhance contact and hence transmission of their diversity and host association patterns.

### (4) Predictors of symbiont sharing in European ants

We hypothesized that the shared evolutionary history of related ant species would promote the sharing of similar symbiont communities. The positive correlation between phylogenetic relatedness of the hosts and symbiont sharing was demonstrated in previous studies on orchid mycorrhiza (Jacquemyn *et al*., 2011) and bat viruses (Luis *et al*., 2015), but no such relationship was found in arbuscular mycorrhiza (Veresoglou & Rillig, 2014) and primate parasites (Cooper *et al*., 2012). Consistent with our prediction, we found that the main factor that promoted symbiont sharing in European ants was the relatedness of the hosts. It indicates that many symbionts pass more easily to related host species. As related ant species employ nearly identical defence structures (nestmate recognition cues, physiological and behavioural defence), it enables symbionts, especially specialized parasites, to by-pass the host defence systems of related hosts. Another key factor that may facilitate the cross-species transmission of symbionts is the overlap in geographical distribution of the hosts (cf. bat viruses in Luis *et al*., 2015). We showed that ant species living in the same biogeographical region possessed more similar symbiont communities. This suggest that both the spatial overlap and similarity in climatic conditions facilitate the sharing of symbionts. Sampling effort also considerably explained the sharing of symbionts. More shared symbionts were reported in well-studied pairs of species. Focusing on the different subsets of ant symbionts, we found that the sharing of both trophobionts and social parasites between host ant species was mainly determined by the same factors, i.e. sample effort, biogeography and phylogenetic relatedness, that drive the sharing of the symbiont community as a whole. However, phylogenetic relatedness was much more important in explaining the sharing of social parasites. This effect is directly linked to the very low taxonomic/phylogenetic distance between their hosts. Myrmecophiles as a whole and the myrmecophilous beetle and mite communities were more similar in sympatric ant species and ant species residing in the same biogeographical region. Interestingly, similarity in these symbiont communities was not positively correlated with host relatedness (cf. Cooper *et al*., 2012; Veresoglou & Rillig, 2014). The host range of myrmecophilous mites is clearly understudied, which is visible in the relative importance of sample effort in the degree of mite sharing.

## V. OUTSTANDING QUESTIONS

Merging different interaction types into one ecological network framework is a key challenge in ecology (Fontaine *et al*., 2011). Diverse host-symbiont communities provide an opportunity to test the relative contributions of ecology and evolution to network assembly. For example, our study on ant-symbiont networks reveal the different role of ecological and evolutionary processes depending on the type of symbiosis. Our insights may provide a basis for theory development and across-ecosystem comparisons (e.g., against plant and coral-based networks) and synthesis.

We lack theory on how the architecture and the interaction signs and sizes jointly affect the stability and productivity of these diverse networks, much in contrast to trophic or mutualistic networks. The relative ease with which one can manipulate ant-symbiont communities makes them suited as empirical systems to test theory.

Host-symbiont networks offer an opportunity to understand both ecological and evolutionary processes behind community assembly, from meso- to macro-ecological scale (see Vellend, 2016). More specifically, as hosts occur spatially structured at these scales, it remains an open question how these assembly processes are determined by ecological and evolutionary dispersal limitation. One key-question here is whether and how symbionts are dispersing: to what degree is horizontal transfer and subsequent symbiont sharing across hosts a facilitator of symbiont community assembly, and to which degree is an obligatory co-dispersal established across the antagonism-mutualism gradient of host-symbiont networks. Mutualistic plant mycorrhizal fungi and plant diaspores are for instance passively co-dispersed by birds (Correia *et al*., 2019). Are alike active processes equally prevalent in ant-symbiont interactions, for instance by symbionts transported by their host during colony relocation (Parmentier, 2019)?

Different host species can coexist locally. In addition to stabilizing and equalizing mechanisms such as competition and niche differentiation, symbiont interactions may directly mediate coexistence of host species.

Insights from this review are restricted and applicable to networks as characterised at the species-level, thereby neglecting any intraspecific variation. Following the relevance of ecological and evolutionary determinants, the question remains open to which degree co-evolutionary dynamics between hosts and their symbiont community occur. As strong selection may act on ant symbionts to bypass host colony defence, cryptic speciation in ant symbionts is expected to be high (Schonrogge *et al*., 2002; Zagaja & Staniec, 2015; von Beeren, Maruyama, & Kronauer, 2015). Symbiont populations may be adapted to an individual host population as was demonstrated in the ant-parasitic syrphid fly *Microdon* and the butterfly *Maculinea* (Elmes *et al*., 1999; Tartally *et al*., 2019). Ultimately, population divergence may result in cryptic symbiont species each targeting a different host species.

At a higher phylogenetic level, other Hymenopteran and insect lineages (Isoptera) provide similar niches to nest symbionts. Our review learned that none of the listed ant symbionts are shared with wasps, solitary and eusocial bees and termites (note that the latter two groups are poorly represented in Europe). Apparently only facultative symbionts (e.g., *Porcellio scaber* in wasp and bee nests for instance) are shared, but more study is needed to understand the drivers of host-symbiont divergence at these deep phylogenetic levels.

## VI. CONCLUSIONS

1. Ant-symbiont networks are particularly interesting to study large-scales patterns and drivers in host-symbiont network topology and symbiont richness as they are extremely diverse and cover the entire mutualism-antagonism continuum. We assembled a complete network of ant-symbiont interactions in Europe and comparatively studied the drivers of host specificity, symbiont richness and symbiont sharing in the different interaction sub-networks.
2. We identified 701 obligate ant macrosymbionts which we categorized in five types: (1) trophobionts: mutualistic aphids and scale insects (*N =* 69) (2) myrmecophiles: commensalistic and parasitic arthropods (*N* = 537), (3) social parasites: parasitic ant species (*N* = 71), (4) ectoparasitic fungi (*N* = 13), and (5) endoparasitic nematodes (*N* = 11).
3. The different types of ant symbionts significantly varied in host specificity. Apart from quantitative differences in host range, we also found clear differences in the average taxonomic/phylogenetic relatedness of the targeted host species for the different types of ant symbionts. The species-richest and best studied ant genera generally supported the largest number of symbionts, but the different types of symbionts were unevenly distributed across the ant genera.
4. We revealed that the ecological and evolutionary factors which drive symbiont species richness may shift depending on the type of symbiosis. Myrmecophile species richness is mainly determined by ecological drivers, such as colony size, host range and niche width of the host. In contrast, species richness of trophobionts and social parasites is driven by evolutionary factors, such as host phylogeny and host biogeography.
5. Ants living in the same biogeographic region shared more symbionts. The sharing of trophobionts and social parasites, in particular, was also strongly facilitated in phylogenetic related hosts.

## VII. SUPPORTING INFORMATION

**Appendix S1.** Host-symbiont matrix listing the associations between ants and ant-symbionts in Europe.

**Appendix S2.** List of literature used to reconstruct the host-symbiont matrix in Appendix S1.

**Fig. S1.** Correlation between the number of described European species in an ant genus and the number of supported symbionts.

**Fig. S2.** Number of host species per symbiont type. Box plots displaying the variation in the number of targeted host species within and across different types of ant symbionts. The width of the boxes is proportional to the square root of reported species in each symbiont group. The different myrmecophilous arthropod orders are displayed in grey.

**Fig. S3.** Phylogenetic distance ± SE, based on the phylogenetic tree of Arnan *et al*. (2017), of targeted host species for different types of ant symbionts. The different myrmecophilous arthropod orders are displayed in grey. Letter codes refer to Tukey post-hoc test. Species with no letters in common are significant different P < 0.05.

## VIII. ACKNOWLEDGEMENTS

This work was supported by Bijzonder Onderzoekfonds Ugent (BOF17/PDO/084 to TP) and FWO (grant nr. 1203020N to TP). DB & FdL are supported by the FWO research network EVENET (W0.003.16N).

## REFERENCES

Aguilée, R., Gascuel, F., Lambert, A. & Ferriere, R. (2018). Clade diversification dynamics and the biotic and abiotic controls of speciation and extinction rates. Nature Communications 9, 1–13.

Antwiki (2019). AntWiki. http://www.antwiki.org/wiki/ [accessed 20 July 2012].

Arnan, X., Cerdá, X. & Retana, J. (2017). Relationships among taxonomic, functional, and phylogenetic ant diversity across the biogeographic regions of Europe. Ecography 40, 448–457.

Barabás, G., D’andrea, R. & Stump, S.M. (2018). Chesson’s coexistence theory. Ecological Monographs 88, 277–303.

Bascompte, J. & Jordano, P. (2007). Plant-animal mutualistic networks: The architecture of biodiversity. *Annual Review of Ecology*, Evolution, and Systematics 38, 567–593.

Bascompte, J., Jordano, P. & Olesen, J.M. (2006). Asymmetric coevolutionary networks facilitate biodiversity maintenance. Science 312, 431–433.

Bastolla, U., Fortuna, M.A., Pascual-García, A., Ferrera, A., Luque, B. & Bascompte, J. (2009). The architecture of mutualistic networks minimizes competition and increases biodiversity. Nature 458, 1018–1020.

Blüthgen, N., Menzel, F., Hovestadt, T., Fiala, B. & Blüthgen, N. (2007) Specialization, constraints, and conflicting interests in mutualistic networks. Current Biology 17, 341– 346.

Boulay, R., Aron, S., Cerdá, X., Doums, C., Graham, P., Hefetz, A. & Monnin, T. (2017). Social life in arid environments: The case study of *Cataglyphis* ants. Annual Review of Entomology 62, 305–321.

Bronstein, J.L., Alarcón, R. & Geber, M. (2006). The evolution of plant-insect mutualisms. New Phytologist 172, 412–428.

Buschinger, A. (2009). Social parasitism among antslj: a review (Hymenopteralj: Formicidae). Myrmecological News 12, 219–235.

Cagnolo, L. & Tavella, J. (2015). The network structure of myrmecophilic interactions. Ecological Entomology 40, 553–561.

Cooper, N., Griffin, R., Franz, M., Omotayo, M. & Nunn, C.L. (2012). Phylogenetic host specificity and understanding parasite sharing in primates. Ecology Letters 15, 1370– 1377.

Correia, M., Heleno, R., Da, Silva, L. P., Costa, J. M. & Rodríguez-Echeverría, S. (2019), First evidence for the joint dispersal of mycorrhizal fungi and plant diaspores by birds. New Phytologist Trust 222: 1054–1060.

Darwin, C. (1859). On the Origin of the Species, or the Preservation of Favoured Races in the Struggle for Life. John Murray, London, UK.

Dáttilo, W., Guimarães, P.R. & Izzo, T.J. (2013). Spatial structure of ant-plant mutualistic networks. Oikos 122, 1643–1648.

Delmas, E., Besson, M., Brice, M.H., Burkle, L.A., Dalla Riva, G. V., Fortin, M.J., Gravel, D., Guimarães, P.R., Hembry, D.H., Newman, E.A., Olesen, J.M., Pires, M.M., Yeakel, J.D. & Poisot, T. (2019). Analysing ecological networks of species interactions. Biological Reviews 94, 16–36.

Donisthorpe, H.S.J.K. (1927). The guests of British ants, their habits and life-histories. George Routledge and Sons, London.

Edwards, D. & Malcolm, F. (2016). Host Specificity among *Unionicola* spp. (Acarilj: Unionicolidae) parasitizing freshwater mussels. The Journal of Parasitology 92, 977– 983.

Elizalde, L., Patrock, R.J.W., Disney, R.H.L. & Folgarait, P.J. (2018). Spatial and temporal variation in host–parasitoid interactions: leafcutter ant hosts and their phorid parasitoids. Ecological Entomology 43, 114–125.

Elmes, G.W., Barr, B., Thomas, J.A. & Clarke, R.T. (1999). Extreme host specifcity by *Microdon mutabilis* (Dipteralj: Syrphidae), a social parasite of ants. Proceedings of the Royal Society London B 266, 447–453.

Evans, G.O. & Till, W.M. (1966). Studies on the British Dermanyssidae (Acari: Mesostigmata) Part II. Classification. Bulletin of The British Museum (Natural History) Zoology 14, 107–370.

Ezenwa, V.O., Price, S.A., Altizer, S., Vitone, N.D. & Cook, K.C. (2006). Host traits and parasite species richness in even and odd-toed hoofed mammals, *Artiodactyla* and *Perissodactyla*. Oikos 115, 526–536.

Feliu, C., Renaud, F., Catzeflis, F., Hugot, J.P., Durand, P. & Morand, S. (1997). A comparative analysis of parasite species richness of Iberian rodents. Parasitology 115, 453–466.

Fontaine, C., Guimarães, P.R., Kéfi, S., Loeuille, N., Memmott, J., Van Der Putten, W.H., Van Veen, F.J.F. & Thébault, E. (2011). The ecological and evolutionary implications of merging different types of networks. Ecology Letters 14, 1170–1181.

Freckleton, R.P., Harvey, P.H. & Pagel, M. (2002). Phylogenetic analysis and comparative data: a test and review of evidence. The American Naturalist 160, 712–726.

Guégan, J., Lambert, A., Lévêque, C., Combes, C. & Euzet, L. (1992). Can host body size explain the parasite species richness in tropical freshwater fishes? Oecologia 90, 197– 204.

Glasier, J.R.N., Poore, A.G.B. & Eldridge, D.J. (2018). Do mutualistic associations have broader host ranges than neutral or antagonistic associations? A test using myrmecophiles as model organisms. Insectes Sociaux 65, 639–648.

Guimarães, P.R., Rico-Gray, V., Dos Reis, S.F. & Thompson, J.N. (2006). Asymmetries in specialization in ant-plant mutualistic networks. Proceedings of the Royal Society B: Biological Sciences 273, 2041–2047.

Hairston, N.G., Smith, F.E. & Slobodkin, L.B. (1960). Community structure, population control, and competition. The American Society of Naturalists 94, 421–425.

Himler, A.G., Adachi-Hagimori, T., Bergen, J.E., Kozuch, A., Kelly, S.E., Tabashnik, B.E., Chiel, E., Duckworth, V.E., Dennehy, T.J., Zchori-Fein, E. & Hunter, M.S. (2011). Rapid spread of a bacterial symbiont in an invasive whitefly is driven by fitness benefits and female bias. Science 332, 254–256.

Hölldobler, B. & Wilson, E.O. (1990). The ants. Harvard University Press, Cambridge, Massachusetts.

Hughes, D.P., Pierce, N.E. & Boomsma, J.J. (2008). Social insect symbionts: evolution in homeostatic fortresses. Trends in Ecology and Evolution 23, 672–677.

Ings, T.C., Montoya, J.M., Bascompte, J., Blüthgen, N., Brown, L., Dormann, C.F., Edwards, F., Figueroa, D., Jacob, U., Jones, J.I., Lauridsen, R.B., Ledger, M.E., Lewis, H.M., Olesen, J.M., Van Veen, F.J.F., Warren, P.H. & Woodward, G. (2009). Ecological networks - Beyond food webs. Journal of Animal Ecology 78, 253–269.

Ivens, A.B.F.,von Beeren, C., Blüthgen, N. & Kronauer, D.J.C. (2016). Studying the complex communities of ants and their symbionts using ecological network analysis. Annual Review of Entomology 61, 353–371.

Ivens, A.B.F., Gadau, A., Kiers, E.T. & Kronauer, D.J.C. (2018). Can social partnerships influence the microbiome? Insights from ant farmers and their trophobiont mutualists. Molecular Ecology 27, 1898–1914.

Jacquemyn, H., Honnay, O., Merckx, V., Brys, R., Tyteca, D., Cammue, B.P.A. & Lievens, B. (2011). Analysis of network architecture reveals phylogenetic constraints on mycorrhizal specificity in the genus *Orchis* (Orchidaceae). New Phytologist 192, 518– 528.

Jaenike, J., Unckless, R., Cockburn, S.N., Boelio, L.M. & Perlman, S.J. (2010). Adaptation via symbiosis: Recent spread of a drosophila defensive symbiont. Science 329, 212–215.

Janet, C. (1897). Etudes sur les fourmis, les guêpes et les abeilles. Note 14: Rapports des animaux myrmécophiles avec les fourmis. Ducourtieux, Limoges.

Kamiya, T., Dwyer, K.O., Nakagawa, S. & Poulin, R. (2014). What determines species richness of parasitic organisms? A meta-analysis across animal, plant and fungal hosts. Biological reviews 89, 123–134.

Kaspari, M. & Vargo, E.L. (1995). Colony size as a buffer against seasonality: Bergmann’s rule in social insects. The American Naturalist 145, 610–632.

Kawakita, A., Okamoto, T., Goto, R. & Kato, M. (2010). Mutualism favours higher host specificity than does antagonism in plant-herbivore interaction. Proceedings of the Royal Society B: Biological Sciences 277, 2765–2774.

Kistner, D.H. (1979). Social and evolutionary significance of social insect symbionts. In Social Insects, Volume I (ed H.R. Herman), pp. 339–413. Academic Press.

Kistner, D.H. (1982). The social insects’ bestiary. In Social Insects, Vol III (ed H.R. Herman), pp. 1–244. Academic Press.

Kronauer, D.J.C. & Pierce, N.E. (2011). Myrmecophiles. Current biology 21, 208–209.

Kuris, A.M., Blaustein, A.R. & Alio, J.J. (1980). Hosts as islands. The American naturalist 116, 570–586.

Leibold, M.A., Holyoak, M., Mouquet, N., Amarasekare, P., Chase, J.M., Hoopes, M.F., Holt, R.D., Shurin, J.B., Law, R., Tilman, D., Loreau, M. & Gonzalez, A. (2004). The metacommunity concept: A framework for multi-scale community ecology. Ecology Letters 7, 601–613.

Lichstein, J.W. (2007). Multiple regression on distance matriceslj: a multivariate spatial analysis tool. Plant Ecology 188, 117–131.

Lindenfors, P., Nunn, C.L., Jones, K.E., Cunningham, A.A., Sechrest, W. & Gittleman, J.L. (2007). Parasite species richness in carnivores: Effects of host body mass, latitude, geographical range and population density. Global Ecology and Biogeography 16, 496– 509.

Luis, A.D., O’shea, T.J., Hayman, D.T.S., Wood, J.L.N., Cunningham, A.A., Gilbert, A.T., Mills, J.N. & Webb, C.T. (2015). Network analysis of host-virus communities in bats and rodents reveals determinants of cross-species transmission. Ecology Letters 18, 1153–1162.

Macarthur, R.H. & Wilson, E.O. (1967). The theory of island biogeography. Princeton University Press.

Martiny, J.B.H., Eisen, J.A., Penn, K., Allison, S.D. & Horner-Devine, M.C. (2011). Drivers of bacterial β -diversity depend on spatial scale. Pnas 108, 7850–7854.

Molero-Baltanás, R., Bach De Roca, C., Tinaut, A., Pérez, J.D. & Gaju-Ricart, M. (2017). Symbiotic relationships between silverfish (Zygentoma: Lepismatidae, Nicoletiidae) and ants (Hymenoptera: Formicidae) in the Western Palaearctic. A quantitative analysis of data from Spain. Myrmecological News 24, 107–122.

Mougi, A. & Kondoh, M. (2012). Diversity of interaction types and ecological community Stability. Science 337, 349–351.

Nava, S. & Guglielmone, A.A. (2013). A meta-analysis of host specificity in Neotropical hard ticks (Acari: Ixodidae). Bulletin of Entomological Research 103, 216–224.

Newman, M.E.J. (2003). The structure and function of complex networks. Siam Review 45, 167–256.

Nosil, P. (2002). Transition rates between specialization and generalization in phytophagous insects. Evolution 56, 1701–1706.

Novgorodova, T.A. (2005). Red wood ants (Formicidae) impact on multi-species complexes of aphids (Aphididae) in the forest-park zone of Novosibirsk. Euroasian Entomological Journal 4, 117–120.

Nunn, C.L., Altizer, S., Jones, K.E., Sechrest, W., Nunn, C.L., Altizer, S., Jones, K.E. & Sechrest, W. (2003). Comparative tests of parasite species richness in primates. The American naturalist 162, 597–614.

Olesen, J.M., Bascompte, J., Dupont, Y.L. & Jordano, P. (2007). The modularity of pollination networks. Proceedings of the National Academy of Sciences of the United States of America 104, 19891–19896.

Pagel, M. (1999). Inferring the historical patterns of biological evolution. Nature 401, 877– 884.

Paracer, S. & Ahmadjian, V. (2000). Symbiosis: an introduction to biological associations. Oxford University Press, New York.

Parmentier, T. (2019). Host following of an ant associate during nest relocation. Insectes Sociaux 66, 329–333.

Parmentier, T. (in press). Guests of social insects. In Encyclopedia of Social Insects (ed C.K. Starr). Springer.

Parmentier, T., Bouillon, S., Dekoninck, W. & Wenseleers, T. (2016). Trophic interactions in an ant nest microcosm: a combined experimental and stable isotope (δ13C/δ15N) approach. Oikos 125, 1182–1192.

Parmentier, T., Dekoninck, W. & Wenseleers, T. (2017). Arthropods associate with their red wood ant host without matching nestmate recognition cues. Journal of Chemical Ecology 43.

Parmentier, T., De Laender, F., Wenseleers, T. & Bonte, D. (2018). Prudent behavior rather than chemical deception enables a parasite to exploit its ant host. Behavioral Ecology 29, 1225–1233.

Pedersen, A.B., Altizer, S., Poss, M., Cunningham, A.A. & Nunn, C.L. (2005). Patterns of host specificity and transmission among parasites of wild primates. International Journal for Parasitology 35, 647–657.

Pimm, S.L. (1979). Complexity and stabilitylj: another look at MacArthur’s original hypothesis. Oikos 33, 351–357.

Poulin, R. (1995). Phylogeny, ecology, and the richness of parasite communities in vertebrates. Ecological Monographs 65, 283–302.

Poulin, R. & Mouillot, D. (2003). Parasite specialization from a phylogenetic perspective: A new index of host specificity. Parasitology 126, 473–480.

Rettenmeyer, C.W., Rettenmeyer, M.E., Joseph, J. & Berghoff, S.M. (2010). The largest animal association centered on one species: the army ant *Eciton burchellii* and its more than 300 associates. Insectes Sociaux 58, 281–292.

Rosas-Valdez, R. & De León, G.P.-P. (2011). Patterns of host specificity among the helminth parasite fauna of freshwater siluriforms: Testing the biogeographical core parasite fauna hypothesis. Journal of Parasitology 97, 361–363.

Sanders, I.R. (2003). Preference, specificity and cheating in the arbuscular mycorrhizal symbiosis. Trends in Plant Science 8, 143–145.

Schoener, T.W. (1989). Food webs from the small to the large: the Robert H. MacArthur award lecture. Ecology 70, 1559–1589.

Schönrogge, K., Barr, B., Wardlaw, J., Napper, E., Gardner, M., Breen, J., Elmes, G. & Thomas, J.A. (2002). When rare species become endangered: Cryptic speciation in myrmecophilous hoverflies. Journal of the Linnean Society 75, 291–300.

Seifert, B. (2007). Die Ameisen Mittel- und Nordeuropas. lutra Verlags- und Vertriebsgesellschaft, Tauer.

Solé, R. V. & Montoya, J.M. (2001). Complexity and fragility in ecological networks. Proceedings of the Royal Society B: Biological Sciences 268, 2039–2045.

Solé, R. V & Bascompte, J. (2006). Self-Organization of Complex Ecosystems (Mpb-42). Princeton University Press.

Stephens, P.R., Altizer, S., Smith, K.F., Alonso Aguirre, A., Brown, J.H., Budischak, S.A., Byers, J.E., Dallas, T.A., Jonathan Davies, T., Drake, J.M., Ezenwa, V.O., Farrell, M.J., Gittleman, J.L., Han, B.A., Huang, S., Et Al. (2016). The macroecology of infectious diseases: a new perspective on global-scale drivers of pathogen distributions and impacts. Ecology letters 19, 1159–1171.

Stoeffler, M., Tolasch, T. & Steidle, J.L.M. (2011). Three beetles—three concepts. Different defensive strategies of congeneric myrmecophilous beetles. Behavioral Ecology and Sociobiology 65, 1605–1613.

Tartally, A., Thomas, J.A., Anton, C., Balletto, E., Barbero, F., Bonelli, S., Bräu, M., Casacci, L. Pietro, Csósz, S., Czekes, Z., Dolek, M., Dziekanska, I., Elmes, G., Fürst, M.A., Glinka, U., Et Al. (2019). Patterns of host use by brood parasitic *Maculinea* butterflies across Europe. Proceedings of the Royal Society B 374, 20180202.

Thomas, J.A., Schönrogge, K. & Elmes, G.W. (2005). Specializations and host associations of social parasites of ants. In Insect Evolutionary Ecology (eds M.D.E. Fellowes, G.J. Hollo & J. Rolff), pp. 479–518. Royal Entomological Society, Cabi Publishing, UK.

Thrall, P.H., Hochberg, M.E., Burdon, J.J. & Bever, J.D. (2007). Coevolution of symbiotic mutualists and parasites in a community context. Trends in Ecology and Evolution 22, 120–126.

Tilman, D. (1982). Resource competition and community structure. Princeton University Press.

Trøjelsgaard, K. & Olesen, J.M. (2013). Macroecology of pollination networks. Global Ecology and Biogeography 22, 149–162.

Tykarski, P. (2017). Coleoptera Poloniae - Information System about Beetles of Poland. http://coleoptera.ksib.pl/?l=en.

Uppstrom, K.A. (2010). Mites (Acari) associated with the ants (Formicidae) of Ohio and the harvester Ant, Messor pergandei, of Arizona. Master thesis, Ohio State University.

Vázquez, D.P., Chacoff, N.P. & Cagnolo, L. (2009). Evaluating multiple determinants of the structure of plant-animal mutualistic networks. Ecology 90, 2039–2046.

Vellend, M. (2016). The Theory of Ecological Communities (Mpb-57), Vol. 75. Princeton University Press.

Veresoglou, S.D. & Rillig, M.C. (2014). Do closely related plants host similar arbuscular mycorrhizal fungal communitieslj? A meta-analysis. Plant and Soil 377, 395–406.

Villa, S.M., Le Bohec, C., Koop, J.A.H., Proctor, H.C. & Clayton, D.H. (2013). Diversity of feather mites (Acari: Astigmata) on Darwin’s finches. Journal of Parasitology 99, 756–762.

Von Beeren, C., Maruyama, M. & Kronauer, D.J.C. (2015). Cryptic diversity, high host specificity and reproductive synchronization in army antassociated *Vatesus* beetles. Molecular Ecology 25, 990–1005.

Wagner, K., Mendieta-Leiva, G. & Zotz, G. (2015). Host specificity in vascular epiphytes: A review of methodology, empirical evidence and potential mechanisms. AoB Plants 7, 1–25.

Wasmann, E. (1894). Kritisches Verzeichniss der myrmekophilen und termitophilen Arthropoden. Berlin: F. L. Dames, xv.

Zagaja, M. & Staniec, B. (2015). *Thiasophila szujeckii* sp. n. (Coleoptera, Staphylinidae, Aleocharinae) - a cryptic species associated with *Formica truncorum* in Poland. Zootaxa 3955, 417–426.

